# Identification of microRNA-target interactions for antioxidant defense response under drought stress by high-throughput sequencing in *Zanthoxylum bungeanum* Maxim

**DOI:** 10.1101/637538

**Authors:** Xitong Fei, Haichao Hu, Jingmiao Li, Yulin Liu, Anzhi Wei

## Abstract

When the plant is in an unfavorable environment such as drought or high temperature, it will accumulate a large amount of active oxygen, which will seriously affect the normal growth and development of the plant. The antioxidant system can remove the reactive oxygen species produced under drought conditions and so mitigate oxidative damage. We examined the trends of antioxidant enzymes, miRNAs and their target genes in Zanthoxylum bungeanum under drought stress. According to the changes of antioxidant enzymes, miRNAs and their target genes expression patterns of *Zanthoxylum bungeanum* under drought stress, an interaction model was constructed to provide a reference for further understanding of plant antioxidant mechanism. The results indicate that under drought stress, POD, CAT, APX, proline, MDA and related genes all show positive responses to drought, while SOD and its genes showed a negative response. It is speculated that in the antioxidant process of *Zanthoxylum bungeanum*, POD, CAT, and APX play a major role, and SOD plays a supporting role. In addition, the expression levels of miRANs and their target genes were basically negatively correlated, indicating that miRNAs are involved in the regulation of the antioxidant system of *Zanthoxylum bungeanum*.

## INTRODUCTION

*Zanthoxylum bungeanum* Maxim. (common name Chinese prickly ash, family *Rutaceae*) is widely distributed in Asia (Yang et al., 2013) where it is an important economic crop. Evolution and natural selection have led the epidermis of *Z. bungeanum* to bear prickles. This species also has strong drought adaptability. The skin of *Z. bungeanum* is the source of one of the eight traditional Chinese condiments, so this plant plays a very important role in Chinese food culture. Because of its unique numbing taste, *Z. bungeanum* is difficult to replace with other seasonings (Zhang et al., 2014). It has become an important component of the diet in various parts of Asia, especially in China. *Zanthoxylum bungeanum* and pepper become best companions and are together an important part of the ‘hot pot’ culture. In addition to its use as a food seasoning, the skin of *Z. bungeanum* also contains chemical components showing proven medicinal properties, including bactericidal (Zhang et al., 2016b), insecticidal (Zhang et al., 2016a), antioxidant (Zhang et al., 2014) and topical anesthetic (Rong et al., 2016).

Drought stress can cause a series of physiological and molecular reactions in plants, which seriously affect normal growth. Thus, drought can cause imbalances in cellular reactive oxygen species (ROS), it can also upset cell membrane lipid peroxidation and it can damage cell and organelle membranes. Excessive ROS have toxic effects on plants. Irrigated agriculture is not yet general, so drought remains one of the most important unfavorable factors affecting both the yield and quality of most commercial crops. Plants have many protective responses to maintain metabolic stability and so continue life under environmental stress. Their antioxidant systems are able to produce a variety of antioxidant enzymes - including superoxide dismutase, peroxidase, antioxidant enzymes, ascorbate peroxidase - to combat the ROS produced under drought stress (Gill & Tuteja, 2010, N. & K., 1977). For example, catalase can decompose H_2_O_2_ produced in plants to form water and oxygen, reducing or eliminating damage by this ROS (Chance. & Maehly., 1955). The antioxidant system also maintains organelle stability, preventing damage to the chloroplast membrane and so stabilizing the PSII system (Lima et al., 2018). In addition, stomata are important gas exchange organs of plants, playing irreplaceable roles in the regulation of photosynthesis, respiration, transpiration and temperature (Li et al., 2017, Martin-StPaul et al., 2017). The stomata are also the water-regulating organs of plants. Under drought, water conservation becomes the decisive factor for plant survival. The response of stomata to drought is also a way for plants to protect themselves. Under drought stress, plants reduce water loss by stomatal closure and so increase their ability to resist drought (Garci’a-Mata. & Lamattina., 2001, Cornic., 2000).

The impact of drought on the yield and quality of *Z. bungeanum* is huge and seriously hinders the development of this industry. miRNAs and their target genes for antioxidant defense response under drought stress remain unclear. Hence a study of their behavior under drought stress, will have significance for better understating this species’ drought-resistance mechanisms. It can also provide a basis for drought-resistant breeding of *Z. bungeanum* and of related species.

## MATERIALS AND METHODS

### Materials

The *Z. bungeanum* seeds were collected from the Fengxian Chinese prickly ash Experimental Station of Northwest Agriculture and Forestry University. They were germinated and cultured in an artificial climate chamber at 25±2°C. Air humidity was set to 80% and the photoperiod to 16:8 h (light:dark). Three-month-old seedlings were used as experimental material. *Zanthoxylum bungeanum* seedlings were then cultured in half-strength Murashige and Skoog (MS) liquid medium containing 20% PEG6000. Leaf samples were collected and stored in liquid nitrogen after periods of 0, 3, 6, 12, 24, 36 and 48 h.

### Methods

#### Physiological index determination

The leaf samples after different periods of drought stress were used to determine antioxidant enzyme activity and malondialdehyde (MDA) and proline contents. Superoxide dismutase (SOD) activity was determined by the nitroblue tetrazolium method (N. & K., 1977). Peroxidase (POD) activity was determined by the guaiacol method and catalase (CAT) activity was determined by the hydrogen peroxide ultraviolet method (Chance. & Maehly., 1955). For APX (L-ascorbate peroxidase) activity we used the method of Panchuk et al (Panchuk et al., 2002). The MDA content was determined by the thiobarbituric acid (TBA) method (Fu. & Huang., 2001). Proline was determined by ninhydrin colorimetry (Bates et al., 1973).

#### Total RNA extraction

Total RNA of the *Z. bungeanum* samples was extracted using the TaKaRa MiniBEST Plant RNA Extraction Kit (TaKaRa, Beijing, China) following the manufacturer’s instructions. The purity and concentration of the RNA obtained were measured using NanoDrop 20000 (Thermo Scientific, Pittsburgh, PA, USA). Only samples where the OD260/280 value was 1.8-2.0 and the OD260/230 value was higher than 2.0 were used for cDNA synthesis.

#### Quantitative Real-time PCR

Primer 7.0 software (Premier, Palo Alto, CA, USA) was used to design RT-qPCT primers (see Table 1). The qRT-PCR assays were carried out on a CFX96 Real-Time PCR Detection System (Bio-Rad, Hercules, CA, USA). The reaction system was of 10 μl, containing 5 μl of 2× SYBR Premix Ex Taq II (TaKaRa), 1 μl of cDNA, 1 μl of each of the upstream and downstream primers and 2 μl of ddH_2_O. *ZbUBQ* and *Zbα-EF* were used for the reference genes to correct the RT-qPCR data (Fei et al., 2018). The RT-qPCT reaction system of miRNAs is identical to mRNA, with a reaction system of 10 μL, containing 5 μL of 2× SYBR Premix Ex Taq II (TaKaRa, Beijing, China), 1 μL of cDNA, 1 μL of each of the forward and universal reverse primer and 2 μL of ddH_2_O. U6 was used for the correction of relative expression levels of miRNAs (Zhang et al., 2018).

**Table 1.**
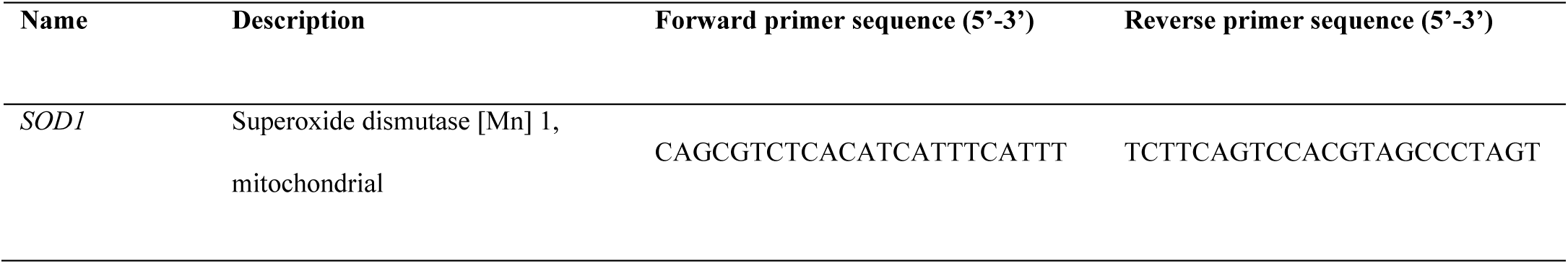
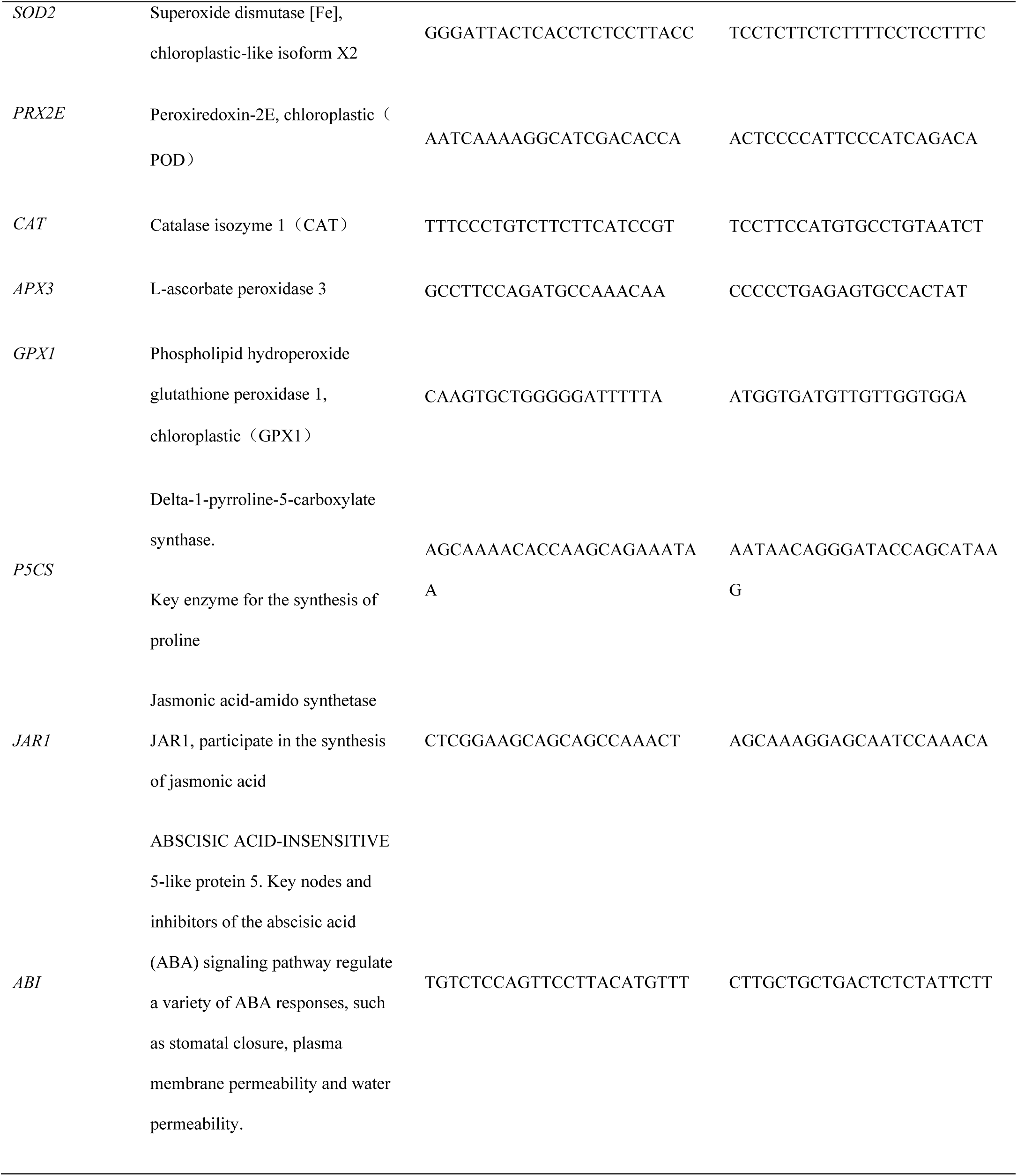
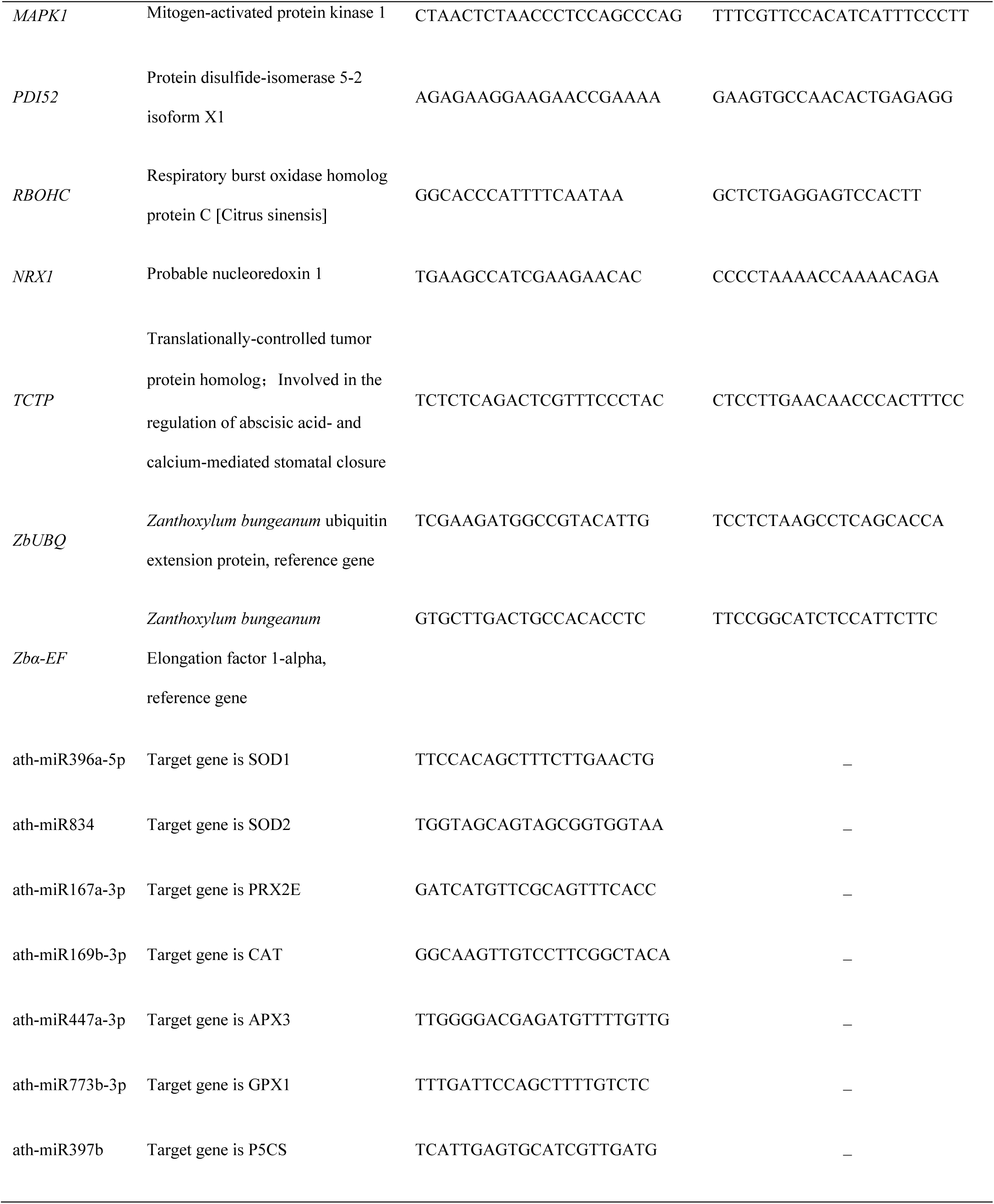
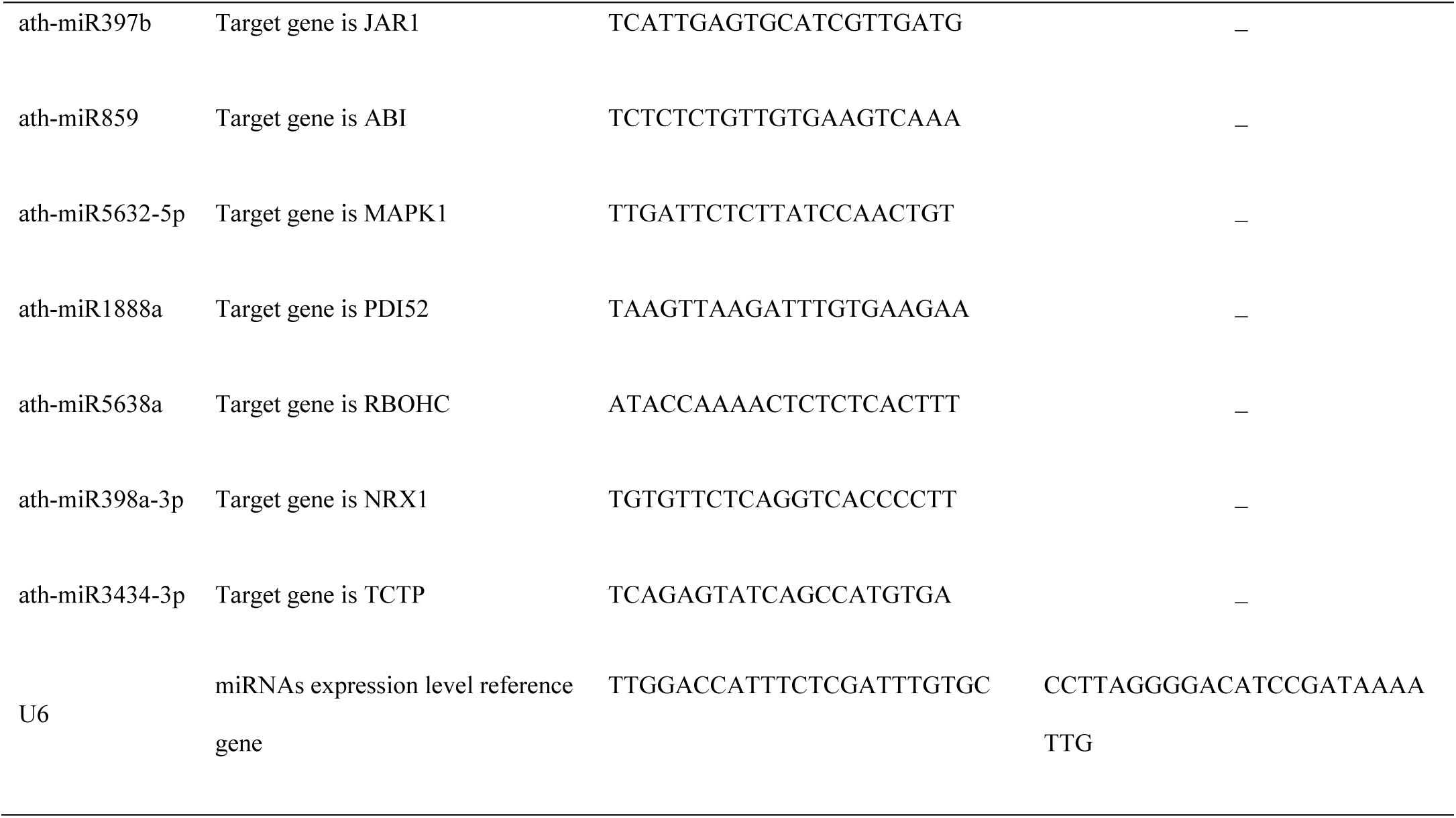
RT-qPCR primers.

#### miRNA prediction

miRNAs that interact with mRNA are predicted in the Arabidopsis miRNA database using the psRNATarget website (http://plantgrn.noble.org/v1_psRNATarget/).

## RESULTS

### Effect of Drought Stress on Antioxidant Enzyme Activity of *Zanthoxylum bungeanum*

Antioxidant enzymes are important roles in plant antioxidant systems. They can eliminate reactive oxygen species in plants and avoid damage to plant plasma membranes. They are also important indicators for evaluating plant antioxidant capacity. The activities of four antioxidant enzymes, SOD, POD, CAT and APX, were determined under drought stress for 7 periods. The results showed that the activity of SOD decreased slowly with the prolongation of drought stress, while POD, CAT and APX decreased. It rose slowly within 6 hours of drought stress, and then increased rapidly between 6h and 24h, and remained at a higher activity level after 24h (Figure 1).

**Figure 1.**
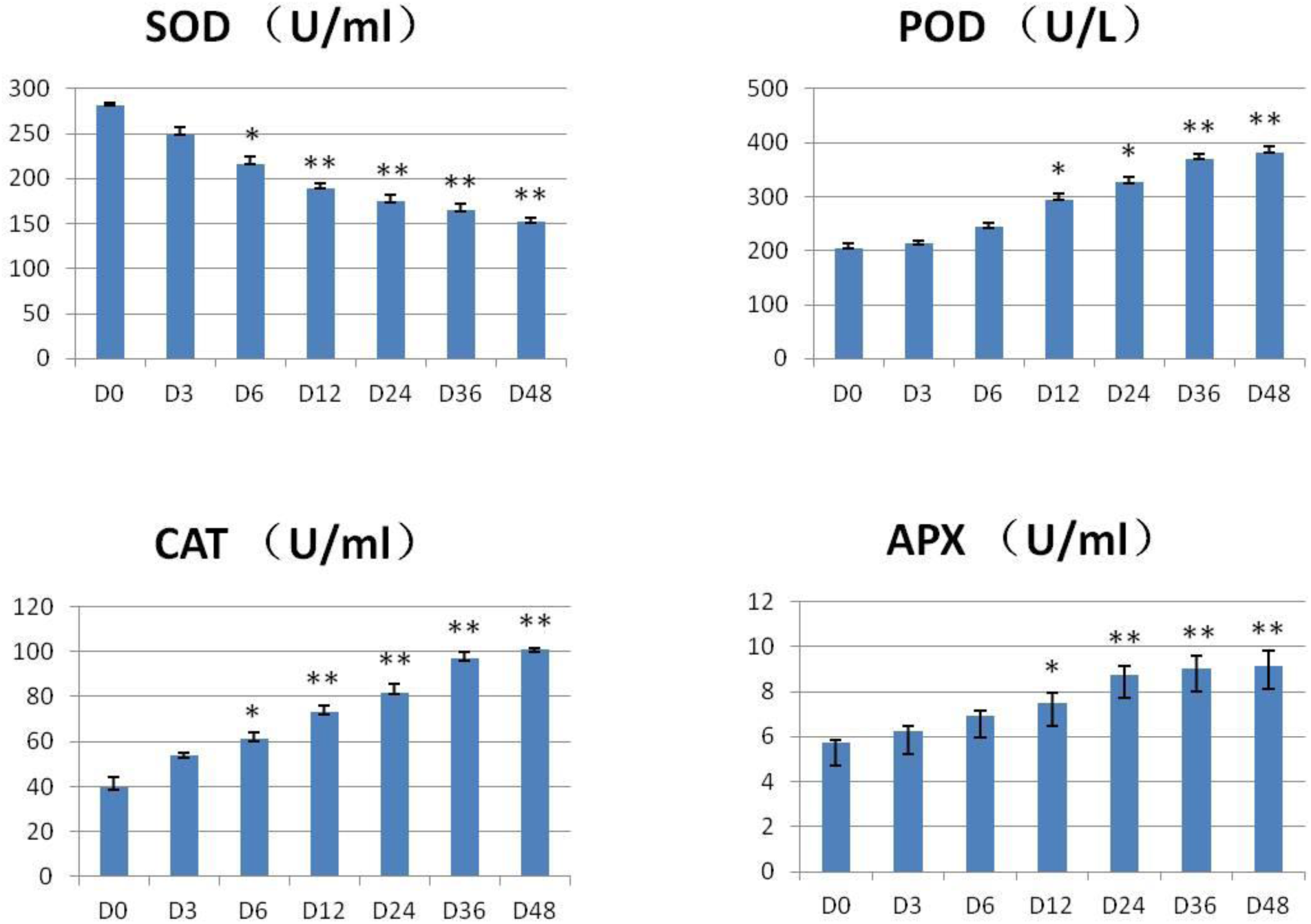
Antioxidant system enzyme activity. * P<0.05; ** P<0.01

The changes in the activity of the four enzymes under drought stress indicate that the response of antioxidant enzymes to drought requires reaction time, and it may be necessary to reach a certain amount of signal to activate the activities of antioxidant enzymes and synthetic pathways. In addition, once activated, antioxidant enzymes are not endlessly synthesized, but remain in a range after a period of drought stress.

Proline is one of the important protective substances for plants to resist stress. Usually, when plants are subjected to abiotic stress, they will synthesize proline in large quantities to cope with adverse environment. Malondialdehyde (MDA) is produced by the peroxidation of membrane lipids in tissues or organs of plants when they are aging or under drought. Therefore, the content of proline and MDA is an important indicator for evaluating plant stress resistance.

The proline content in Zanthoxylum bungeanum was gradually increased from 13 μg/ml at the initial stage of drought stress to 20 μg/ml at 48 h. The content of MDA changed more vigorously, and it quickly exceeded 12 nmol/ml within 12 hours, which was more than twice that of D0 period, and then remained above 14 nmol/ml for 24h to 48h (Figure 2). The increase of proline and MDA content indicates that in the process of drought stress of *Zanthoxylum bungeanum*, on the one hand, the peroxidation of organ membrane lipids affects the physiological function of plants, on the other hand, the protective substances are also synthesized to slow down the damage of plants caused by adverse environment.

**Figure 2.**
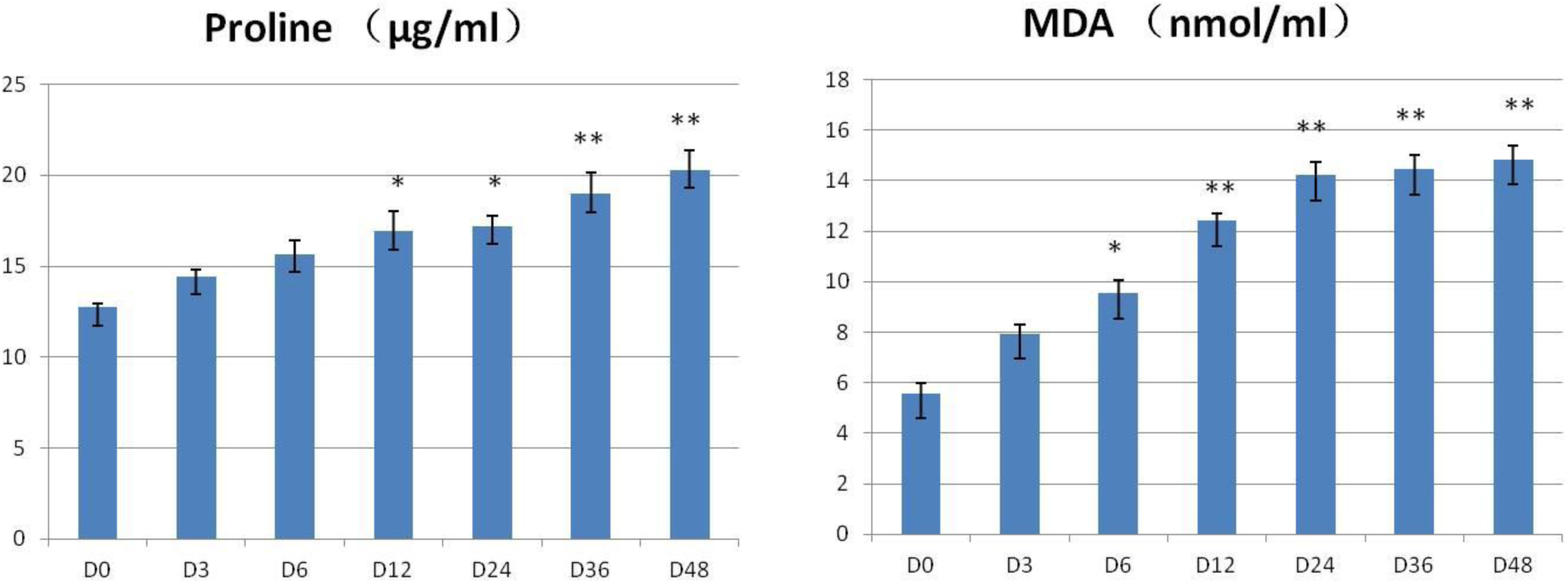
Proline and MDA contents. * P<0.05; ** P<0.01

### Expression pattern of miRNAs and their target genes under drought stress

The expression patterns of genes related to the antioxidant system were essentially consistent with the changes in the related substances. Genes such as *PRX2E, CAT, APX3, P5CS* and *GPX1* showed a significant increase under drought stress (Figure 3).

**Figure 3.**
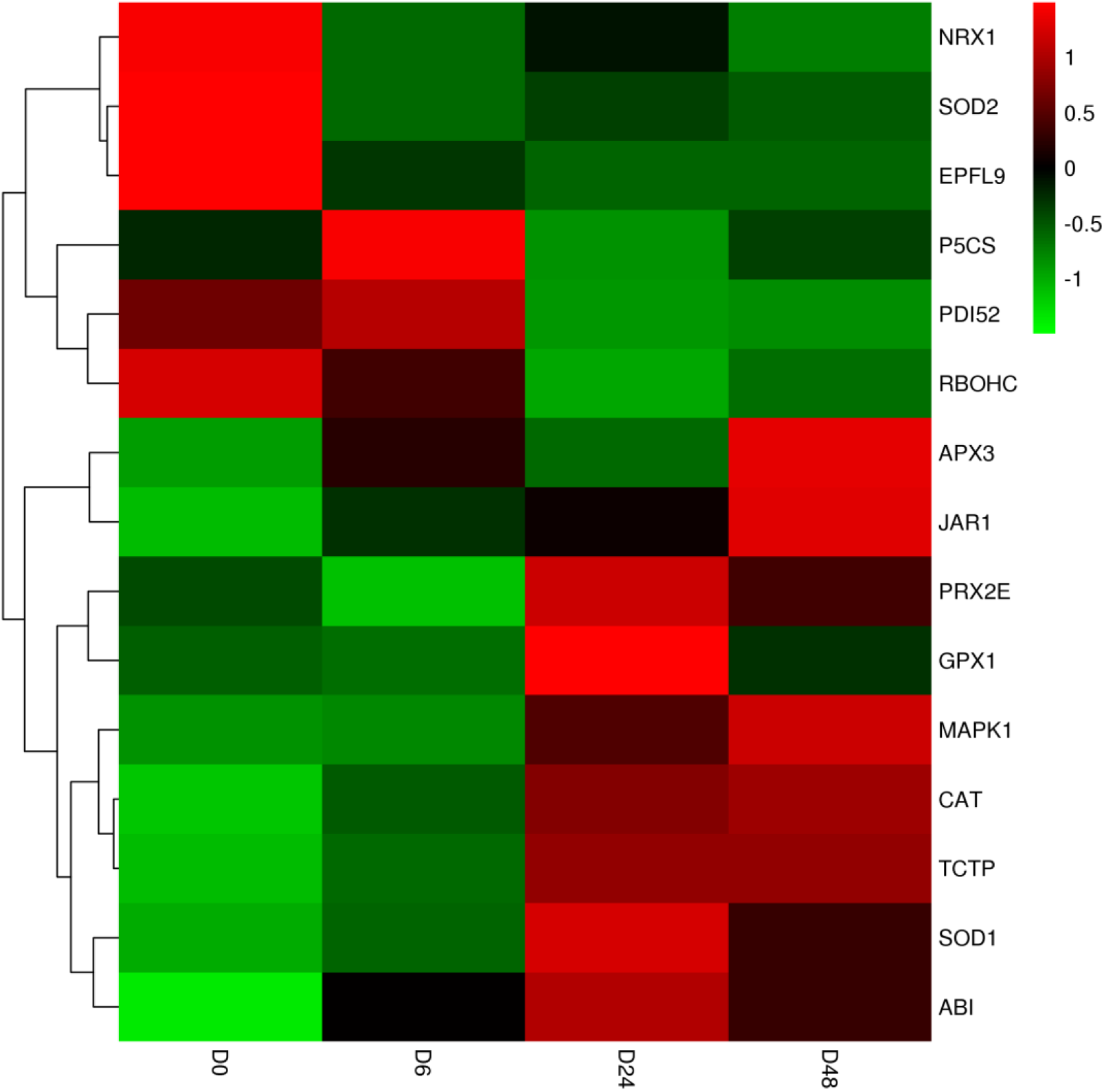
Heat map of gene expressions relating to the antioxidant system.

miRNAs interacting with mRNAs were predicted via the psRNATarget website (Table 2). The predicted results indicate that ath-miR396a-5p binds to *SOD1* and inhibits its transcription. Ath-miR167a-3p interacts with *PRX2E*, and *CAT* is the target gene of ath-miR169b-3p. In addition, we also predicted miRNAs interacting with other genes on the drought stress-related signaling pathway, involving jasmonic acid signaling pathway, ABA signaling pathway, MAPK signaling pathway and proline synthesis pathway. Furthermore, the relative expression levels of miRNAs regulating the genes involved in the synthesis of antioxidant enzymes were detected. The results showed that the expression trends of miRNAs and their target genes were basically negatively correlated (Figure 4). It can be stated that these miRNAs are important regulators involved in the antioxidant system of *Z. bungeanum*.

**Table 2.**
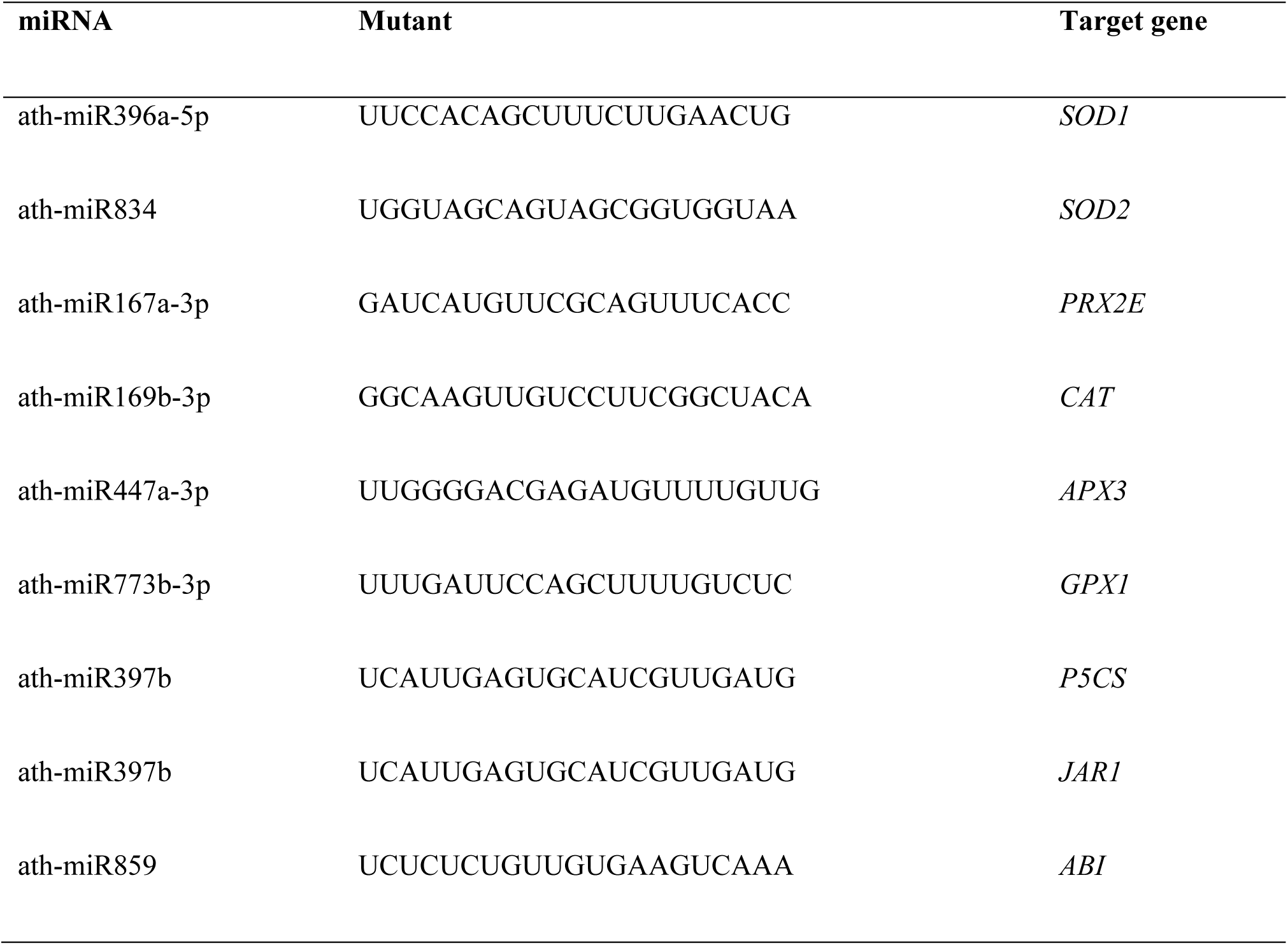
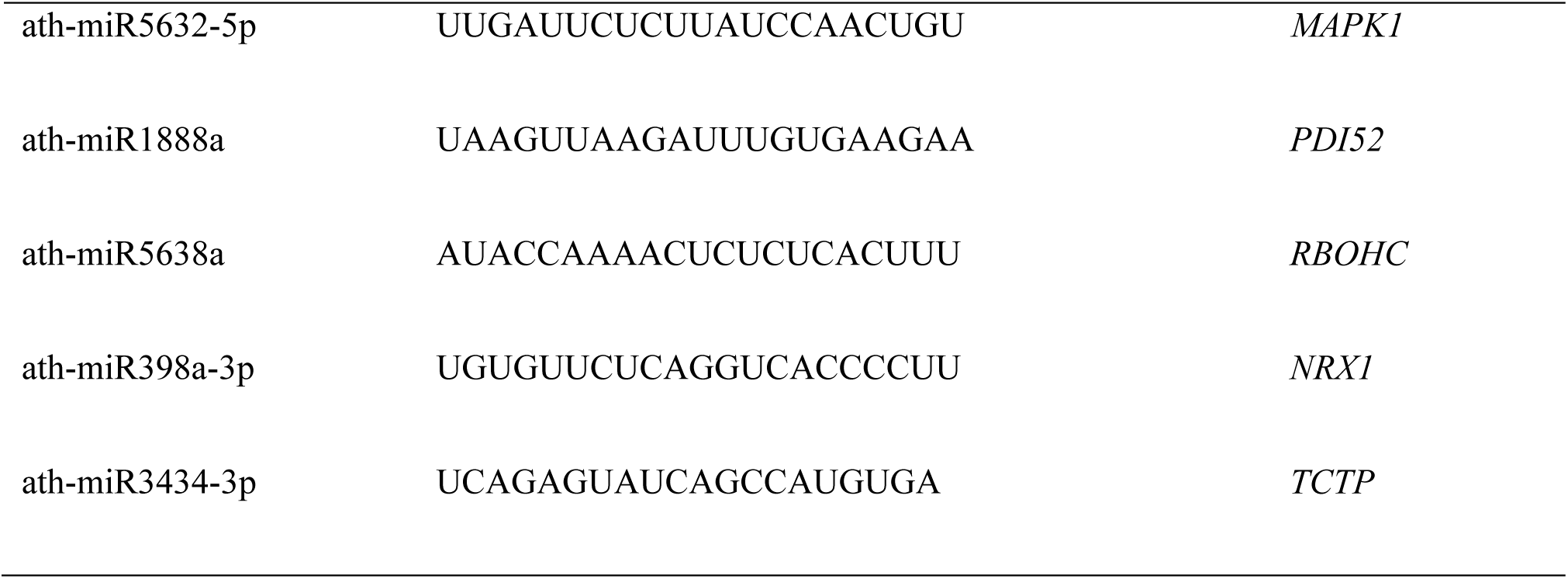
miRNAs and their target genes.

**Figure 4.**
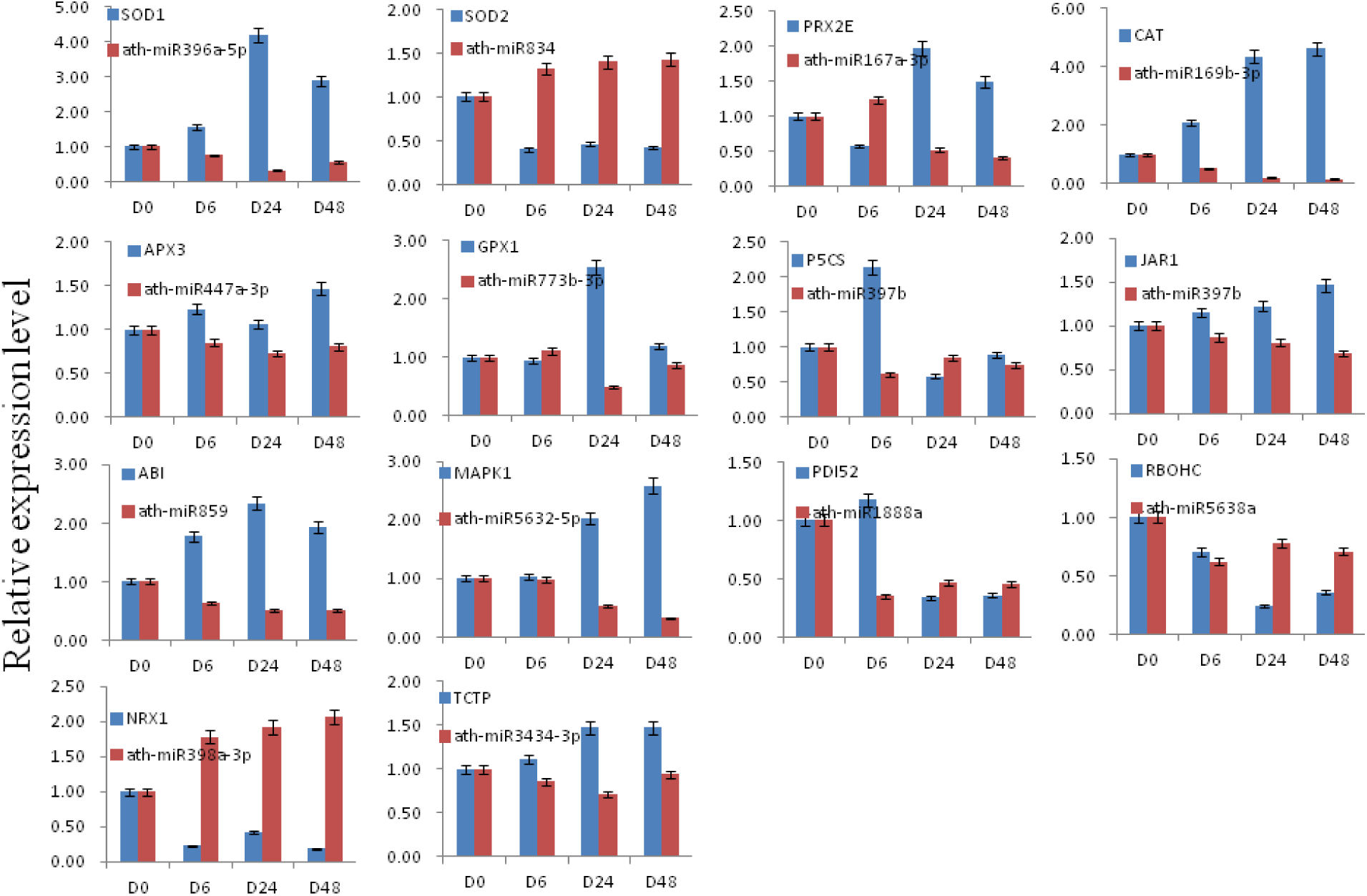
miRNAs and their target genes.

The expression levels of the SOD gene in chloroplasts and mitochondria were monitored and we found the *SOD2* gene in the chloroplast was positively correlated with superoxide dismutase activity. The *SOD1* gene in mitochondria was negatively correlated with the activity of superoxide dismutase. It is concluded that the superoxide dismutase produced by *Z. bungeanum* under drought stress comes mainly from the chloroplast.

In addition, some expression patterns of pathway genes activated by drought stress were also monitored. *P5CS* is a key enzyme in the proline synthesis process. *ABI* is a key inhibitor of the ABA signaling pathway and participates in the closure of stomata and *TCTP* is involved in ABA and calcium ion-mediated stomatal closure. In addition, the relative expression levels of *MAPK1* and *PDI52* were also up-regulated (Figure 4). At the same time, the relative expression of JAR1, a gene related to jasmonic acid synthesis, was also up-regulated under the induction of drought stress. However, other genes were inhibited, such as *RBOHC, NRX1, EPFL9*.

## DISCUSSION

### Response of antioxidant enzymes under drought stress

The results show that most of antioxidant enzymes trend upward under continuing drought stress, while SOD shows a downward trend. The trend of the *SOD2* gene and SOD in the chloroplasts are consistent, which indicates the chloroplast antioxidant system was severely damaged during the drought. In addition, it can be explained that in the antioxidant system, several other antioxidant enzymes (such as POD, CAT, APX) exert major antioxidant effects. Proline and MDA increased gradually during the drought stress and may also be involved in signal transduction and protection. Studies have shown that *Barbula fallax* and *Zanthoxylum bungeanum* have similar patterns of antioxidant enzyme activity, where SOD showed a downward trend in the early stage of drought stress, while POD and CAT activities were positive in response to drought stress and increased in the early stage of drought stress (Zhang et al., 2017). In many species, the activities of antioxidant enzymes such as SOD, POD, CAT, and APX generally rise under drought. Chickpea accumulates proline and increases the activity of SOD, APX, GPX and CAT under drought stress (Dalvi et al., 2017). The activities of SOD, POD and CAT in alfalfa increase significantly under drought stress (Tina et al., 2017). This also occurs in pea (Mittler. & Zilinskas., 1994), rice (Sharma & Dubey, 2005), Kentucky bluegrass (Bian & Jiang, 2009) and sesame (FAZELI. et al., 2007).

### Antioxidant signaling pathway regulatory factors interaction

Many signal pathways are activated in plants under drought stress, including signal transduction, gene interaction, physiological changes and so on. Under drought stress, the accumulation of ROS in plants seriously affects the normal growth and development of plants. Antioxidant system can effectively prevent the damage caused by ROS produced by plants, and is an indispensable self-protection system for plants. It plays a very complex process in the antioxidant system, involving the synthesis of antioxidant enzymes, transcription and modification of functional genes, transport of ions, and so on. According to the experimental results and previous research results, the factors involved in the regulation of antioxidant system were analyzed and a signal regulation model of antioxidant system was constructed.

Under drought, mitochondrial respiration can produce ROS. Oxygen produced by chloroplast photosynthesis and external oxygen are also sources of ROS in plants (Figure 5a). Reactive oxygen species can destroy cell membranes and interfere with normal growth of plants, while the accumulation of ROS is toxic to plants. The production of ROS activates the plant’s antioxidant system. SOD, POD, CAT, APX, GPX and other enzymes can convert superoxide anion to H_2_O_2_ and eventually decompose it into non-toxic H_2_O and O_2_ (Wang et al., 2018, Duan et al., 2009) (Figure 5b).

**Figure 5.**
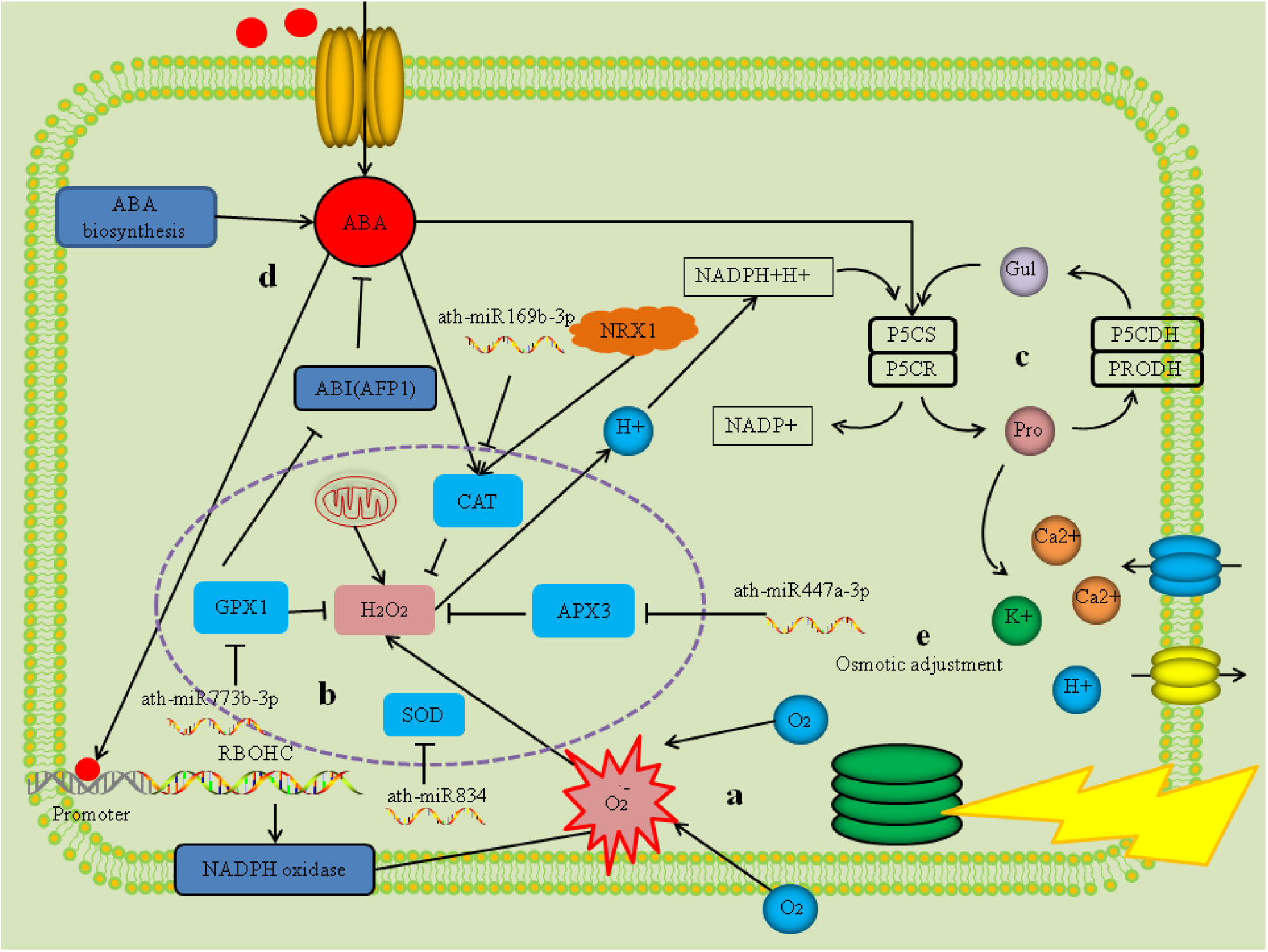
Antioxidant signaling pathway genes interaction model. (a) Oxygen produced by photosynthesis and oxygen in vitro are the main sources of ROS. (b) SOD, POD, CAT, APX, GPX convert superoxide anion to hydrogen peroxide and eventually decompose into oxygen and water. (c) The antioxidative enzyme decomposes H^+^ produced by hydrogen peroxide as a substrate for the synthesis of proline from glutamate. (d) ABA can bind to the *RBOHC* promoter to promote the synthesis of NADPH oxidase. (e) Proline can regulate the concentration of ions in cells.

In this process, miRNAs are involved in the regulation of the synthesis of antioxidant enzymes, directly inhibiting the transcription or degradation of the corresponding mRNA, resulting in a decrease in the amount of antioxidant enzyme synthesis. However, by analyzing the expression levels of miRNAs, the expression levels of miRNAs associated with antioxidant systems are mostly declining. The above analysis can show that in order to protect itself from ROS damage under drought stress, plants have largely relieved the inhibitory effect of miRNAs on antioxidant system. During the scavenging of H_2_O_2_ by antioxidant enzymes, it is susceptible to oxidative stress, resulting in reduced clearance. NRX1 is able to reduce oxidized antioxidant enzymes and has a stable antioxidant system (Kneeshaw. et al., 2017). However, in the gene expression level study, the expression level of *NRX1* was down-regulated. It is concluded that drought interfered with the reduction of NRX1 against oxidase. The antioxidant system decomposes ROS to produce H+, which provides a substrate for glutamate synthesis in the proline pathway (Figure 5c).

Proline is an important osmo-regulatory substance, and the main way to regulate the osmotic potential is to regulate the concentration of ions in the cell. Proline produced under drought stress protects cells from damage by controlling the concentration of ions, and becomes an important regulator of plant self-protection (Tieleman et al., 2001, Woolfson et al., 1991). *P5CS* is an important synthetic gene in the proline synthesis pathway, and ath-miR397b can inhibit the expression level of *P5CS*. Under drought stress, the relative expression level of ath-miR397b was decreased, which inhibited the inhibition of *P5CS* and promoted the synthesis of proline. The accumulation of ROS can activate the Ca^2+^ channel on the cell membrane(Figure 5e), causing a large amount of Ca^2+^ to enter the cell, and so can increase the Ca^2+^ concentration of the guard cells and change their osmotic potential (Singh et al., 2017). At the same time, high concentration of Ca^2+^ can suppress the input of K^+^ and the output of H^+^. In addition, SLAC1 transports anions out (Vahisalu et al., 2008), resulting in an increase in the concentration of cations in the membrane. The combination of ABA and TCTP can induce stomatal closure, and CPK can phosphorylate CAT as well as promote stomatal closure (Zou et al., 2010).

At the same time, plants can also synthesize ABA under drought stress, and ABA can promote the synthesis of proline from glutamate, which eventually leads to a large accumulation of proline (Figure 5d) (Strizhov. et al., 1997). Proline is an important osmotic adjustment substance in plants, so the above reaction is beneficial, allowing plants to cope better with drought. ABA binds to the promoter of RBOHC and promotes the production of respiratory burst oxidase (NADPH oxidase) (Zhao. et al., 2001). NADPH oxidase (NOX) also activates Ca^2+^ channels on the cell membrane (Kurusu et al., 2015). In addition, it has been shown to play an important protective role in plant drought stress, preventing leaves from being destroyed by ROS (Duan et al., 2009, Miller. et al., 2009). Under drought stress, the ABA signaling pathway and antioxidant system of plants are activated, and there is close interaction between them (Qi. et al., 2015, He et al., 2018). On the one hand, ABA can promote the synthesis of CAT and improve the efficiency of the antioxidant system. While, on the other hand, *GPX1* can inhibit *ABI*, thereby relieving the inhibition of ABA by plants (Lim et al., 2017).

## ACKNOWLEDGMENTS

The authors would like to thank Yao Ma for his participation in the manuscript discussion. This study was financially supported by the National Key Research and Development Program Project Funding (2018YFD1000605).

